# Cortical lipids containing choline mediate cannabinoid-induced cognitive improvement

**DOI:** 10.1101/2024.03.07.583670

**Authors:** Marta Moreno-Rodríguez, Jonatan Martínez-Gardeazabal, Iker Bengoetxea de Tena, Alberto Llorente-Ovejero, Laura Lombardero, Estibaliz González de San Román, Lydia Giménez-Llort, Iván Manuel, Rafael Rodríguez-Puertas

## Abstract

Recent research connecting choline-containing lipids to basal forebrain cholinergic neurons (BFCN) degeneration in neuropathological states highlights a challenge for balancing lipid integrity with optimal acetylcholine (ACh) levels. Warranting an adequate choline source to maintain ACh levels in this pathway is crucial for preserving memory. The endocannabinoid (eCB) system plays a role in modulating learning and memory processes controlled by cholinergic neurotransmission. Consequently, we propose that activation of this system is neuroprotective against cholinergic degeneration. In the present study, we investigated the neuroprotective effect of a subchronic treatment with the CB_1_ cannabinoid agonist, WIN55,212-2, using both *ex vivo* and *in vivo* 192IgG-Saporin models of specific cholinergic damage. Degeneration of baso-cortical cholinergic pathways induced memory deficits and a downregulation of saturated and mono-unsaturated lysophosphatidylcholines (LPC) cortical levels. WIN55,212-2 not only restored memory deficits but also increased cortical ACh levels and modified cortical choline-containing lipids such as sphingomyelins (SM) and LPCs, which are essential for correct memory functioning, in lesioned animals. Given these results, we propose that WIN55,212-2 generates an alternative choline source through the breakdown of SMs, which is enough to increase cortical ACh levels and LPCs. These findings suggest that modification of choline-containing lipids by the activation of CB_1_ receptors is a promising therapy for dementia associated with cholinergic dysfunction, such as in Alzheimer’s disease (AD).

## INTRODUCTION

The selective vulnerability of basal forebrain cholinergic neurons (BFCN) plays a crucial role in the pathophysiology of dementia in Alzheimer’s disease (AD)^1, 2^. A significant loss of cholinergic neurons in the nucleus basalis of Meynert and decreased levels of presynaptic cholinergic markers in the neocortex were described, correlating with cognitive decline in AD^3, 4^. Currently, the largest class of drugs approved for the treatment of AD are inhibitors of the enzyme acetylcholinesterase to increase acetylcholine (ACh) at the synaptic cleft, however the clinical benefits of these drugs are limited. Therefore, there is a significant need to develop novel drugs to enhance the functionality of the BFCN projection system especially when damage has already occurred^5^. Recently, studies have successfully traced cholinergic pathways *in vivo*, demonstrating that the integrity of these pathways is disrupted not only in patients with mild cognitive impairment (MCI) and AD, but also in individuals with subjective cognitive decline^6, 7^. Given the importance of the above-described cholinergic neurotransmission in AD, animal models of cholinergic dysfunction based on experimental manipulations of the BFCN have been developed as an appropriate tool to study the memory deficits^8,9^. While the BFCN lesion model does not exhibit the histopathological characteristics of AD such as neurofibrillary tangles and βA plaques, as seen in genetic models of AD like the 3xTg-AD mouse model^10^, it provides a valuable tool for exploring treatments targeted at improving cognition after cholinergic damage has occurred. Cholinergic neurons are unique in their requirement of choline, which is used to synthesize both choline-containing lipids (i.e., phosphatidylcholine (PC), lysophosphatidylcholine (LPC), choline plasmalogen, and sphingomyelin (SM)) and their neurotransmitter, ACh^11, 12^. The recent description of a close association between choline-containing lipids and BFCN degeneration in AD^13^, suggests that, under pathological conditions, the cholinergic system may encounter a dilemma, having to choose between preserving the structural integrity of choline-containing lipids in the membrane and maintaining optimal levels of ACh. Therefore, it is crucial to understand the vulnerabilities of BFCN and explore novel approaches and pathways to maintain cholinergic system integrity.

The endocannabinoid (eCB) system is a neuromodulator system that plays important roles in learning and memory processing, distributed in areas of the brain related to cognition^14^ and implicated in the cholinergic neurotransmission^15, 16^. Cannabinoid agonists induce memory impairment^17, 18^, but in the last decade, evidence has been accumulating showing a beneficial effect of low cannabinoid doses upon cognitive impairment^19–21^. The role of the cannabinoid system in neurodegenerative diseases is still unknown; however, increased activity of cannabinoid receptor 1 (CB_1_) has been observed with disease progression^22^. Additionally, a case report revealed that micro-dosing of cannabinoids improved mnemonic learning in a patient with AD^23^. Although some studies showed that cannabinoids modulated the ACh release in the hippocampus and cortex^24–26^, the specific mechanism through which cannabinoids impact or enhance memory remains unknown.

Consequently, to investigate memory deficits and the role of the eCB system in a model of BFCN degeneration, our group previously employed intraparenquimal injections of the p75^NTR^-binding 192IgG-Saporin toxin into the nucleus basalis magnocellularis (NBM) in rats. The studies showed that after the lesion, rats showed memory impairment, increased levels of CB_1_ receptor activity^27^ and altered levels of choline-containing lipids^28^ in both the NBM and cortex. These results suggested that choline-containing lipids and the eCB system play a key role in the specific degeneration of basal forebrain-cortical cholinergic circuit. These findings led us to employ cannabinoid agonists as a therapeutic approach to treat cholinergic deficits.

In this study, we have used the mentioned in vivo animal model of BFCN degeneration to present evidence of an alternative cellular source of choline that uses the synthetic cannabinoid WIN55,212-2 to restore acetylcholine levels, the choline-containing lipids, to induce cognitive improvement.

## MATERIALS & METHODS

### Animals

*Ex vivo* hemibrain organotypic cultures were derived from 25 male Sprague-Dawley rats postnatal day 7 (P7), weighting 14-20 g, and for the *in vivo* experimental model, 121 adult male Sprague-Dawley rats, weighting 200-250 g, were used for 192IgG-saporin or vehicle administration. All rats were housed in cages (50 cm length x 25 cm width x 15 cm height), four or five per cage, at 22°C in a humidity-controlled (65%) room with a 12:12 hour’s light/dark cycle, with access to food and water *ad libitum*. 7-month-old C57BL/6 3xTg-AD mice (n = 17) harboring PS1M146V, APPSwe, and TauP301L genes provided by Prof. Lydia Giménez-Llort from Universitat Autònoma de Barcelona and age-matched wild-type C57BL/6 (n = 20) from Envigo (Indianapolis, IN, USA) weighing 25-30 g were also used. Mice were housed in groups of 3-4 per cage at a temperature of 22°C and in a humidity-controlled (65%) room with a 12:12 hours light/dark cycle, with access to food and water *ad libitum*. Every effort was made to minimize the discomfort of the animals and to use the minimum number of animals. All procedures were performed in accordance with European animal research laws (Directive 2010/63/EU) and the Spanish National Guidelines for Animal Laws (RD 53/2013, Law 32/2007). Experimental protocols used in this study were approved by the Local Ethics Committee for Animal Research of the University of the Basque Country (CEEA M20-2018-52 and 54).

### *Ex vivo* model of cholinergic degeneration in organotypic cultures and cannabinoid treatments

P7 Sprague-Dawley rats were sacrificed by decapitation and brains were quickly dissected under aseptic conditions inside a laminar flow cabinet. The protocol used was described in detail by Llorente-Ovejero et al.^28^. In brief, approximately 6 slices containing cholinergic neurons within the NBM were obtained from each brain, and these were immediately transferred into cell culture inserts over membranes of 0.4 µm pore size (PIC50ORG, Millipore, MA, USA), placed in 6-well culture dishes (Falcon, BD Biosciences Discovery Labware, Bedford, MA) containing cell culture medium. The culture medium consisted in 49% (v/v) neurobasal medium (NB, Sigma-Aldrich), 24% (v/v) Hanks’ balanced salt solution (HBSS, Gibco), 24% (v/v) normal horse serum (NHS, Gibco), 1% (v/v) d-glucose, 0.5% glutamine (Sigma-Aldrich), 0.5% B27 supplement serum free (Gibco), and 1% antibiotic/antimycotic. The culture plates were incubated at 37°C in a fully humidified atmosphere supplemented with 5% CO_2_. The *ex vivo* hemibrain organotypic cultures were randomly divided into two groups: In group 1 fresh cell culture medium was added. In group 2 fresh cell culture medium containing 192IgG-saporin (Millipore Temecula, CA, USA) (100 ng/ml) was added on days 2 and 5 *in vitro* (DIV). Both groups were treated with WIN55,212-2 (1 nM or 10 nM) (Sigma-Aldrich, St. Louis, MO, USA) dissolved in ethanol. The maximum final ethanol concentration in culture medium was set at 0.01% (v/v), according to previous reports ^29^. To verify receptor specificity of the effect exerted by the cannabinoid agonist WIN55,212-2, a third group of animals were treated as those in group 2 with the addition of CB_1_ receptor antagonist AM251 (1 µM) (Tocris Bioscience, Bristol, UK). After 8 DIV, organotypic cultures were incubated in the presence of 5 µg/ml of propidium iodide (PI) to mark degenerating cells (bright red) for 2 h prior to fixation with paraformaldehyde.

### *In vivo* rat model of basal forebrain cholinergic degeneration and cannabinoid treatments

Basal forebrain cholinergic degeneration was induced following bilateral stereotaxic (−1.5 mm anteroposterior from Bregma, ±3 mm mediolateral from midline, +8 mm dorsoventral from cranial surface) injection of 192IgG-saporin (130 ng/µl) into the NBM, as previously described^27^. Control rats received an injection of artificial cerebrospinal fluid (aCSF) into the NBM. Rats were allowed to recover from surgery for 7 days. On day 8, we initiated treatments and training on the Barnes maze (BM) and novel object recognition test (NORT), as described below.

In the BM performance, aCSF and 192IgG-SAP groups received intraperitoneal (ip) injections of WIN55,212-2 (0.5 mg/kg or 3 mg/kg) (Sigma-Aldrich, St. Louis, MO, USA) or vehicle solution (1:1:18; DMSO:Kolliphor: saline) for five consecutive days, 1 h prior to the performance of the task. To verify receptor specificity of the cannabinoid effect, another group of animals received an ip injection of WIN55,212-2 (0.5 mg/kg) along with CB_1_ receptor antagonist SR141716A (0.5 mg/kg, ip) (Tocris Bioscience, Bristol, UK). Rats were randomly selected for each group: aCSF (n = 8); aCSF +W0.5 (n=7); aCSF +W3 (n=6); aCSF +SR (n=6); 192IgG-SAP (n=8); 192IgG-SAP+W0.5 (n=7); 192IgG-SAP+W3 (n=6); 192IgG-SAP+W+SR (n=6); 192IgG-SAP+SR (n=6).

In the NORT performance, WIN55,212-2 (0.5 mg/kg, ip, 5 days) was administered daily for five consecutive days, 1 h before each phase of the behavioral test. The following groups of rats were used: control group aCSF (n=10), aCS+WIN0.5 (n=10), 192IgG-saporin (n=10), 192IgG-SAP+W0.5 group (n=10). Animals were sacrificed three days after the last WIN55,212-2/SR141716A/vehicle administration.

### WIN55,212-2 administration in the 3xTg-AD mouse model of familial AD

Given that the loss of basal forebrain cholinergic projections is an early feature of AD, we studied if the same cannabinoid treatment would also be beneficial in an animal model of familial AD, the 3xTg-AD mouse, which shows the histopathological hallmarks of the disease^30^. WIN55,212-2 (0.1 mg/kg, equivalent to 0.5 mg/kg in rats, ip, 5 days)^31^ was administered daily for five consecutive days, 1 h before each phase of BM test to 3xTg-AD and age-matched wild-type C57BL/6 mice. The following groups of animals were used: control group (WT, n=10), WIN55,212-2 (0.1 mg/kg) group (WT+WIN0.1, n=10), 3xTg-AD group (3xTg-AD, n=8) and 3xTg-AD + WIN55,212-2 (0.1 mg/kg) group (3xTg-AD+WIN0.1, n=9).

### Barnes maze

The maze was performed using two white circular platforms, one for rats (130 cm of diameter, 100 cm from the floor, 20 holes 10 cm each and 2.5 cm between holes) and one for mice (92 cm of diameter, 100 cm from the floor, 20 holes 5 cm each and 2.5 cm between holes). Only one of the holes leads to a dark chamber located under it. Two bright lights (400 W, approximately 1310 luxes light condition) and visual cues were placed around the platform. Each rodent was placed in the middle of the maze and was allowed to explore the maze for 3 minutes. If a rodent did not reach the target hole in the given 3 minutes, it was gently guided to it. During 4 days of training, rodents conducted 4 trials per day, with 15 minutes between trials. During the training days, total latency (the time to reach the target hole) was measured. A gradual decrease in this parameter over the 4 training days is indicative of spatial memory. On day 5, the target hole was closed and rodents were allowed to explore the maze for 3 minutes. As an additional measure of spatial memory, time in the target quadrant (the quadrant where the target hole was located) was measured. The maze was cleaned using a 10% ethanol solution after every trial. All the procedures were analyzed by SMART 3.0 video tracking software (Panlab Harvard apparatus, Barcelona, Spain).

### Novel object recognition test

The test was performed in a white open-field arena (90 × 90 × 50 cm) (Panlab S.L., Barcelona, Spain) in a room under one lux light condition. A video camera placed above the shuttle box recorded the behavior of the rats. The test was divided into four distinct phases that were carried out throughout five days: habituation phase (3 days), familiarization phase, short-term testing (5 h after familiarization) and long-term testing (24 h after familiarization). Before each phase, rats were transported to the experimental room for about 10 min and each rat was gently handled individually for 1 min, having its neck and back stroked by the experimenter’s fingers, before entering the arena. After leaving the arena, rats were gently handled again. Habituation phase lasted for 3 days and consisted in placing rats in the arena to allow them to explore the compartment for 5 min. In the familiarization phase, which was carried out on the fourth day, rats were presented with two identical objects (object A and object A), built with five to six mega bloks, with a height of about 10 cm. The objects were positioned diagonally in opposite corners of the arena, approximately 10 cm away from their respective walls, and were mirror images of each other. To avoid possible bias regarding the location of the objects, these were rotated after the familiarization phase of each rat. A 25 s exploration threshold for both objects combined was established and rats remained in the arena until that threshold was met. If rats failed to reach the 25 s exploration threshold in 15 min, they were excluded from the study. Exploration of the objects was considered when the rats touched the object or faced it with their nose being less than 2 cm away from it. 5 h after the familiarization phase, short-term testing was performed. In that phase, rats were again placed in the arena and were presented with one of the familiar objects (object A) and with a new object (object B). Rats were given 5 min to explore the objects. 24 h after the familiarization phase, on the fifth day, long-term testing was performed. Rats were again placed in the arena and were presented with the familiar object (object A) and a third, new object (object C). Rats were given 5 min to explore the objects. In the first habituation phase, which is equivalent to an open field test, the total path length of the rats and their speed were measured using an automated tracking system (SMART, Panlab S.L., Barcelona, Spain) as indicators of exploratory behavior. In the short and long-term testing phases, the amount of time dedicated to exploring the familiar and new objects was measured and the object discrimination ratio (DR) was calculated using the following formula: DR = [(Novel Object Exploration Time - Familiar Object Exploration Time) / Total Exploration Time]. A higher DR was indicative of more time exploring the new object compared to the familiar one and was thus considered a positive performance in the test (good recognition memory). DR scores approaching zero reflect no preference for the new object and negative scores indicate preference for the familiar object, which reflect impairment of recognition memory in both cases. Moreover, the total exploration time spent by the rats in short- and long-term testing phases was measured to investigate the effect of the different model or the drugs administered on object exploration.

### Tissue Preparation

Organotypic cultures on day 8 were gently and extensively rinsed with 0.9% saline solution (37°C) followed by immersion in 4% paraformaldehyde and 3% picric acid in 0.1M PB (4°C) for 1 h. Animal groups which performed the BM, on day 15 after the lesion, were anesthetized with ketamine/xylazine (90/10 mg/kg; ip) and sacrificed by decapitation or transcranial perfused to obtain fresh or fixed tissue, respectively. Fresh brains from experimental groups (n=96) were quickly removed by dissection, fresh frozen, and kept at −80 °C. Later, brains were cut into 20 μm coronal sections using a Microm HM550 cryostat (Thermo Scientific, Waltham, MA, USA) equipped with a freezing-sliding microtome at −25 °C and mounted onto gelatin-coated slides and stored at −25 °C until used. Animals from experimental groups (n=25) were transcardially perfused with 50 mL of warm (37 °C), calcium-free Tyrode’s solution (0.15 M NaCl, 5 mM KCl, 1.5 mM MgCl_2_, 1 mM MgSO_4_, 1.5 mM NaH_2_PO_4_, 5.5 mM glucose, 25 mM NaHCO_3_; pH 7.4), 0.5% heparinized, followed by 4% paraformaldehyde and 3% picric acid in 0.1 M phosphate buffer (PB) (4 °C) (100 mL/100 g, bw) (37 °C, pH 7.4). Brains were removed and placed in a cryoprotective solution consisting of 20% sucrose in PB overnight at 4 °C, and frozen by immersion in isopentane and kept at −80 °C. Brains were cut into 12 μm coronal sections as described above, mounted onto gelatin-coated slides and stored at −25 °C until used for the immunofluorescence assays.

### Immunofluorescence

Organotypic culture sections were blocked and permeabilized with 4% normal goat serum (NGS) with 0.6% Triton X-100 in PBS (0.1 M, pH 7.4) for 2 h at 4°C. The incubation was performed using the free-floating method at 4°C (48 h) with rabbit anti-p75^NTR^(1:500; Cell signaling, MA, USA) with 0.6% Triton X-100 in PBS with 5% BSA. The primary antibody was then revealed by incubation for 30 min at 37°C in darkness with donkey anti-rabbit Alexa 488 (1:250; Thermo Scientific, Waltham, MA, USA) with Triton X-100 (0.6%) in PBS. For the processing of fixed rat tissue, 12 μm coronal sections were blocked and permeabilized with 3% donkey serum with 0.25% Triton X-100 PBS (0.1 M, pH 7.4) and 2 h later they were labeled with mouse anti-Iba1 (1:500; Fujifilm Wako Chemicals, VA, USA) or rabbit anti-p75^NTR^ (1:750; Cell signaling, MA, USA) overnight. After several washes, the appropriate secondary antibody (1:200) was applied (Donkey anti-rabbit Alexa fluor-488 for p75^NTR^ and donkey anti-mouse Alexa fluor-555 for Iba1; Thermo Scientific, Waltham, MA, USA) for 2 h. Controls of immunofluorescence consisted in primary antibody omission resulting in the absence of immunoreactivity.

### Cells quantitation

In organotypic cultures, 200-fold magnification photomicrographs of the BFCNs within the NBM were acquired by means of an Axioskop 2 Plus microscope (Zeiss) equipped with a CCD imaging camera (SPOT Flex Shifting Pixel). Both p75^NTR^ immunoreactive and PI positive cells were stereologically counted and the total number of cells in the whole image was reported. The population of p75^NTR^ immunoreactive or PI-stained cells was expressed as p75^NTR^ or PI cells/mm^2^. In rat tissue, 200-fold magnification photomicrographs of NBM in both hemispheres were randomly acquired by Axioskop 2 Plus microscope (Carl Zeiss) equipped with a CCD imaging camera SPOT Flex Shifting Pixel. p75^NTR^ immunoreactive positive cells were stereologically counted at three different stereotaxic levels (-1.20 mm, -1.56 mm and -1.92 mm from Bregma), in 6 different rats per group and the total number of cells in the whole image was obtained. The density of cholinergic cells was expressed as p75^NTR^ positive cells/mm^3^. Iba1 immunoreactive positive cells were counted at stereotaxic level -1.56 mm from Bregma, in 3 different rats per group and the total number of cells in the whole image was obtained. The number of Iba1 positive cells was expressed as cells/mm^2^.

### MALDI-MSI

Matrix-assisted laser desorption/ionization mass spectrometry imaging (MALDI-MSI) was performed using fresh 20 μm sections for each sample. MBT matrix was deposited on the tissue surface by sublimation. The sublimation was performed using 300 mg of MBT, and the deposition time and temperature were controlled (23 min, 100°C). For the recrystallization of the matrix, the sample was attached to the bottom of a glass Petri dish face-down, which was placed on another Petri dish containing a methanol-impregnated piece of filter paper on its base. The Petri dish was then placed on a hot plate (1 min, 38 °C)^32^. A MALDI LTQ-XL-Orbitrap (Thermo Fisher Scientific, San Jose, CA, USA) equipped with a nitrogen laser (λ = 337 nm, rep rate = 60 Hz, spot size = 80 μm × 120 μm) was used for mass analysis. Thermo’s ImageQuest software was used to analyze MALDI-MSI data and image acquisition in positive ion mode. The used range was 400–1000 Da with 10 laser shots per pixel at a laser fluence of 15 μJ. The target plate stepping distance was set at 150 μm for both x- and y-axes by the MSI image acquisition software. The data were normalized using the total ion current values. Each of the m/z values was plotted for signal intensity for each pixel (mass spectrum) across a given area (tissue section) using MSiReader software^33^. The m/z range of interest was normalized using the ratio of the total ion current for each mass spectrum. The data were expressed as absolute intensity in arbitrary units. The assignment of lipid species was facilitated using the databases Lipid MAPS (http://www.lipidmaps.org/) and the Human Metabolome Database (HMDB) (https://hmdb.ca). 5 ppm mass accuracy was selected as the tolerance window for the assignment.

### [^35^S]GTPγS autoradiography

Fresh 20 µm slices from all experimental groups were dried, followed by two consecutive incubations in HEPES-based buffer (Sigma-Aldrich, St. Louis, MO, USA) (50 mM HEPES, 100 mM NaCl, 3 mM MgCl_2_, 0.2 mM EGTA and 0.5% BSA, pH 7.4) for 30 min at 30°C to remove endogenous ligands. Then, slices were incubated for 2 h at 30°C in the same buffer but supplemented with 2 mM GDP, 1 mM DTT, adenosine deaminase (3 Units/l) (Sigma-Aldrich, St. Louis, MO, USA) and 0.04 nM [^35^S]GTPγS (1250 Ci/mmol, PerkinElmer, Boston MA, USA). Basal binding was determined in two consecutive slices in the absence of the agonist. The agonist-stimulated binding was determined in another consecutive slice with the same reaction buffer, but in the presence of the corresponding receptor agonists, CP55,940 (10 µM) (Sigma-Aldrich, St. Louis, MO, USA) for CB_1_ receptor receptors and carbachol (100 µM) for M_2_/M_4_ receptors. Non-specific binding was defined by competition with non-radioactive GTPγS (10 µM) (Sigma-Aldrich, St. Louis, MO, USA) in another section. Then, slices were washed twice in cold (4°C) 50 mM HEPES buffer (pH 7.4), dried and exposed for 48h to β-radiation sensitive film with a set of [^14^C] standards (American Radiolabeled Chemicals, St. Louis, MO, USA) calibrated for [^35^S].

### Histochemistry for AChE detection

Fresh 20 µm slices from all experimental groups were air dried and post-fixed with 4% paraformaldehyde for 30 min at 4°C. Slices were rinsed twice in 0.1 M Tris-maleate buffer (pH 6.0) for 10 min and incubated in the AChE reaction buffer: 0.1 M Tris-maleate; 5 mM sodium citrate; 3 mM CuSO_4_; 0.1 mM iso-OMPA; 0.5 mM K_3_Fe(CN)_6_ and 2 mM acetylthiocholine iodide (Sigma-Aldrich, St. Louis, MO, USA) as reaction substrate. The incubation time to stain cholinergic fibers was 100 min. The enzymatic reaction was stopped by two consecutive washes (2x10 min) in 0.1 M Tris-maleate (pH 6.0). Slices were then dehydrated in increasing concentrations of ethanol and covered with DPX as the mounting medium. Finally, the stained slices were scanned at 600 ppi resolution, the images were converted to 8-bit gray-scale mode and AChE positive fiber density was quantified by Image J software (NIH, Bethesda, MD, USA). Software measured the optical density (O.D.) of AChE reactivity in each anatomical area.

### Choline/Acetylcholine assay

Choline and acetylcholine were quantified in rat cortical tissue from all experimental groups using a choline/acetylcholine assay kit (ab65345, Abcam, Cambrige, UK). 10 mg of cortical fresh tissue were harvested, washed in cold PBS and resuspended in 500 μL of choline assay buffer. The tissue was homogenized with a homogenizer (Heidolph RZR 50 Homogenizer 300-2000 RPM w/ Barnant 50001-92 Stand 115V), sitting on ice, with 10 – 15 passes. Samples were centrifuged at 4°C for 5 minutes (Eppendorf 5417R Refrigerated Centrifuge) at 16,400 rpm. Supernatants were collected and used with the choline/acetylcholine assay buffer. The assay was carried out in accordance with the manufacturer’s instructions, in the absence and presence of acetylcholinesterase to identify values of total and free choline, which allowed an indirect quantification of acetylcholine. The relative sample fluorescence was determined using Varioskan LUX Reader (Thermo Scientific, Waltham, MA, USA).

### Rat brain cortex incubation

10 mg of fresh cortical tissue were harvested, washed in cold PBS and resuspended in 500 μL of choline assay buffer. The tissue was homogenized with a homogenizer (Heidolph RZR 50 Homogenizer 300-2000 RPM w/ Barnant 50001-92 Stand 115V), sitting on ice, with 10 – 15 passes. The samples were incubated in choline assay buffer at 37 °C and collected every 15 minutes, for 2 h. After incubation, samples were centrifuged at 4°C for 5 minutes (Eppendorf 5417R Refrigerated Centrifuge) at 16,400 rpm. Supernatants were collected to use them with the choline/acetylcholine assay buffer and dry pellets were analyzed by MALDI-MS.

### Cortical sample preparation for MALDI-MS

Cortical lipid composition was analyzed in all experimental groups by MALDI-MS. Dry pellets from rat cortical tissue from all experimental groups were obtained and the protein concentration was determined using the Bradford method. Samples were reconstituted with water at the same concentration. A mixed sample (3 μL of sample and 7 μL of matrix-saturated solution of MBT) was deposited on a MALDI plate containing 96 wells, using the dried droplet method. Xcalibur software was used for MALDI data acquisition in both positive and negative ion modes. The positive ion range was 400–1000 Da, and the negative ion range was 400–1100 Da, with 3 minutes of shots per well at a laser fluence of 15 μJ. The m/z range of interest was normalized using the ratio of the total ion current for each mass spectrum. The data were expressed as absolute intensity in arbitrary units. The assignment of lipid species was facilitated using the databases Lipid MAPS (http://www.lipidmaps.org/) and the Human Metabolome Database (HMDB) (https://hmdb.ca). 5 ppm mass accuracy was selected as the tolerance window for the assignment.

### Statistical analysis

Data are expressed as mean ± SEM. Data evaluated across the groups used Kruskal-Wallis test followed by Dunn’s *post hoc* tests for multiple comparisons. Spearman rank for correlations (SigmaPlot 12.5) and false discovery rate were used to adjust for multiple comparisons between correlations. Statistical significance was set at *p* < 0.05 (two-tailed). Statistics and data were graphically represented using GraphPad Prism 9 (GraphPad Software). The heat map generation was performed using the freely available software program Heatmapper (http://www.heatmapper.ca/).

## RESULTS

### WIN55,212-2 protects cell viability after BFCN degeneration in organotypic culture

Propidium iodide (PI) uptake was quantified as a measure of cell death in NBM after 192IgG-saporin and different cannabinoid treatments in rat postnatal day 7 (P7) hemibrain organotypic cultures. *Ex vivo* culture application of 100 ng/ml of the toxin 192IgG-saporin at days two and five produced a significant increase in the density of PI-stained cells (PI^+^ cells /mm^2^; Control: 14.22 ± 2 *vs* 192IgG-saporin: 63.3 ± 7, \*\**p<*0.01. Supplementary Figure 1 A, C). Pre-treatment of organotypic cultures with either 1 nM or 10 nM of WIN55,212-2, 2h prior to the application of 192IgG-saporin, induced protective effects on cell viability (192IgG-SAP: 63.3 ± 7 *vs* 192IgG-SAP+W [10 nM]: 17.78 ± 7, ^#^*p*<0.05. Supplementary Figure 1 A, C). p75^NTR+^ cells in the same area were counted as specific marker of cholinergic neuron. The application of the immunotoxin at days two and five led to a statistically significant decrease in the density of cholinergic cells in NBM (p75^NTR+^ cells /mm^2^; aCSF: 61.87 ± 4 *vs* 192IgG-SAP: 26.22 ± 7, \**p*<0.05. Supplementary Figure 1 B, C), while pre-treatment of cultures with both doses of WIN55,212-2, 2h prior to the application of the toxin, did not change p75^NTR+^ cells number. Although WIN55,212-2 did not specifically protect cholinergic cell viability, it induced a secondary protective effect in the viability of the whole cell population of the organotypic cultures.

### WIN55,212-2 restored spatial and recognition memory on an *in vivo* model of cholinergic degeneration

Rats received bilateral intraparenchymal injections of 192IgG-saporin into the region containing the NBM and were subsequently tested for memory performance on the BM and NORT tests. As WIN55,212-2 did not provide protection to cholinergic neurons in the *ex vivo* experiments, a single dose of WIN55,212-2 was administered for five consecutive days starting on day eight once the lesion had stabilized.

In the BM, a progressive reduction in the total latency of the experimental groups indicates proper memory function during the acquisition training (Figure 1 A). In probe day, lesioned animals spent significantly less time in the target quadrant compared to control group (Time in the target quadrant (s); aCSF: 91.41 ± 3 *vs* 192IgG-SAP: 49.22 ± 3, \*\*\**p*<0.001, Figure 1 B and C), showing memory impairment after toxin administration. The administration of both doses of WIN55,212-2 (0.5 and 3 mg/kg) after the BFCN lesion increased the time in the target quadrant, reaching control levels (aCSF: 91.41 ± 3, 192IgG-SAP+W0.5: 86.82 ± 5, 192IgG-SAP+W3: 90.64 ± 9, Figure 1 C). Co-treatment with SR141617A, a specific CB_1_ receptor antagonist, blocked WIN55,212-2 cognitive improvement, indicating that cognitive restoration was mediated by the activation of CB_1_ receptor. Interestingly, the high dose of WIN55,212-2 produced opposite effects: cognitive function was improved in lesioned rats, while it was impaired in control rats (Figure 1 C). The findings of a reduced time spent in the target quadrant of control group following the administration of 3 mg/kg of WIN55,212-2 supports the classic detrimental effects of cannabinoid agonists on memory, while the beneficial effect of both doses on lesioned rats suggests a biphasic effect of cannabinoids on cognition.

**Figure 1.**
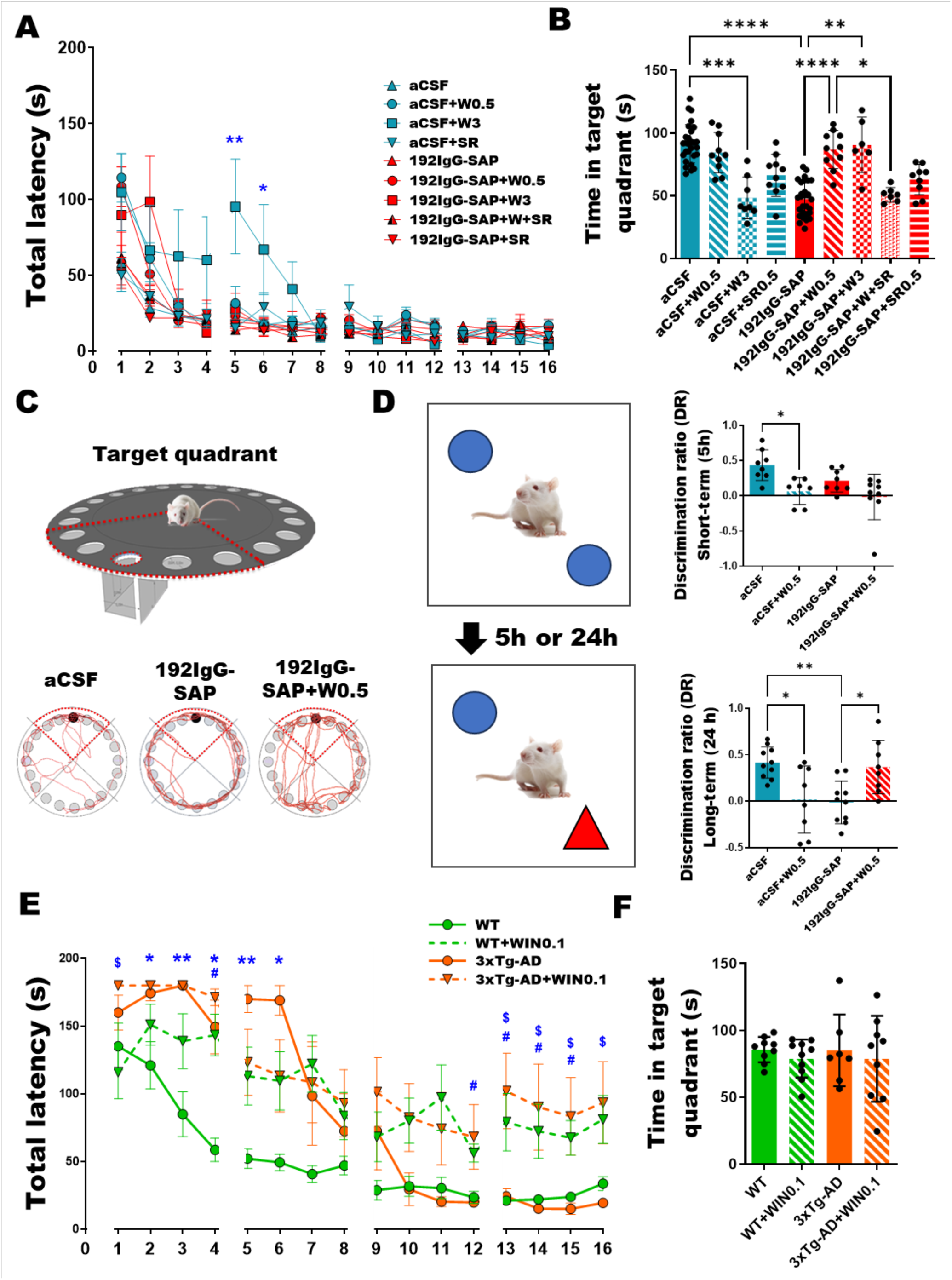
WIN55,212-2 improved cognitive impairment evaluated by BM and NORT following BFCN lesion. (A). Analysis of the total latency, which is the time spent by the rats to reach the target hole during 16 trials throughout 4 days for all the groups (B). Time in target quadrant of the rats on day 5, which is the time spent in the target quadrant. aCSF, aCSF+W0.5, 192IgG-SAP+W0.5 and 192IgG-SAP+W3 spent more time in target quadrant than the rest of experimental groups. (C) Image of the Barnes Maze with the target quadrant delineated in red and the trajectories of aCSF, 192IgG-SAP and 192IgG-SAP+W0.5 groups. (D) On the left a scheme depicting a simplified version of the protocol followed for the performance of NORT. On the right total exploration time of the objects in the short-term and the long-term for aCSF, aCSF +W0.5, 192IgG-SAP and 192IgG-SAP+W0.5 (E) The total latency of WT, WT+WIN0.1, 3xTg-AD and 3xTg-AD+WIN0.1 groups (F) Analysis of the time spent in the target quadrant of the mice from WT, WT+WIN0.1, 3xTg-AD and 3xTg-AD+WIN0.1 groups in BM test on probe trial day. No significant differences were observed between the groups, indicating that all four groups performed well in the test on probe trial day. (Kruskal–Wallis test, *post-hoc* test Dunn’s multiple comparison \*\*\**p*<0.001).

To complete this study, we employed the most effective dose from the BM, 0.5 mg/kg, to analyze the discrimination ratio in NORT in both the short-term (5 h post-familiarization with the objects) and the long-term (24 h post-familiarization). In the short-term, the subchronic treatment with WIN55,212-2 impaired memory in control rats (Figure 1 D), as already observed with the high dose 3 mg/kg in the BM. The short-term recognition memory was relatively preserved in 192IgG-SAP group. In the long-term test, WIN55,212-2 clearly impaired recognition memory in control rats as measured by a decrease in the DR, while 192IgG-SAP caused a significant decrease in the DR in the long-term (aCSF 0.42 ± 0.05 *vs* 192IgG-SAP 0.06 ± 0.07, *p*< 0.01; Figure 1 D), indicating recognition memory impairment following the depletion of BFCNs in the long-term, but not in the short-term. Importantly, the administration of WIN55,212-2 to lesion rats improved memory in NORT test in the long-term, increasing the DR to control levels (SAP 0.06 ± 0.07 *vs* SAP+WIN0.5 0.38 ± 0.13, *p*< 0.05; Figure 1 D), consistent with the observations in the BM test and further indicating a different effect of cannabinoid agonists on memory depending on the cognitive status of the subject.

Finally, we investigated whether the same cannabinoid treatment would also yield benefits in an animal model of familial AD, the 3xTg-AD mouse, in the BM. WIN55,212-2 was administered to both wild-type (WT) and 3xTg-AD mice at a dose of 0.1 mg/kg, equivalent to 0.5 mg/kg in rats^31^. The time to reach the target hole during each trial showed significant differences between the groups. On day 4 of the acquisition phase, both WT and 3xTg-AD mice showed reduced total latencies, indicating a correct learning process for both phenotypes, which was significantly slower for 3xTg-AD mice, suggesting mild spatial cognitive deficits at 7 months of age in this AD model. WIN55,212-2 administration in the WT induces a deleterious effect on learning, while in 3xTg-AD mice, the treatment did not reverse the observed cognitive deficits (Figure 1 E). In the probe trial, which is the greatest indicator of positive performance in this test, no statistically significant differences were observed between groups, suggesting an absence of spatial memory deficits in the four groups (Figure 1 F). Consequently, we decided to focus the study in the rat model of BFCN degeneration.

### WIN55,212-2 did not modify the glial response following the 192IgG-saporin administration

Glial activation following a lesion of the NBM was analyzed in aCSF, 192IgG-SAP, aCSF+W0.5 and 192IgG-SAP+W0.5 groups. Following 192IgG-saporin administration, quantification revealed that the number of p75^NTR+^ cells decreased significantly (p75^NTR^ cells/mm^3^; aCSF: 786 ± 42 *vs* 192IgG-SAP: 123 ± 30, \*\*\**p*<0.001, Figure 2 A, C), while the number of Iba1 positive cells increased (Iba1 cells/mm^2^; aCSF: 477 ± 41 *vs* 192IgG-SAP: 636 ± 30, \**p*<0.05. Figure 2 B, C). After WIN55,212-2 administration, the number of p75^NTR^ or Iba1 positive cells at the lesion site were not modified (Figure 2 A-C).

**Figure 2.**
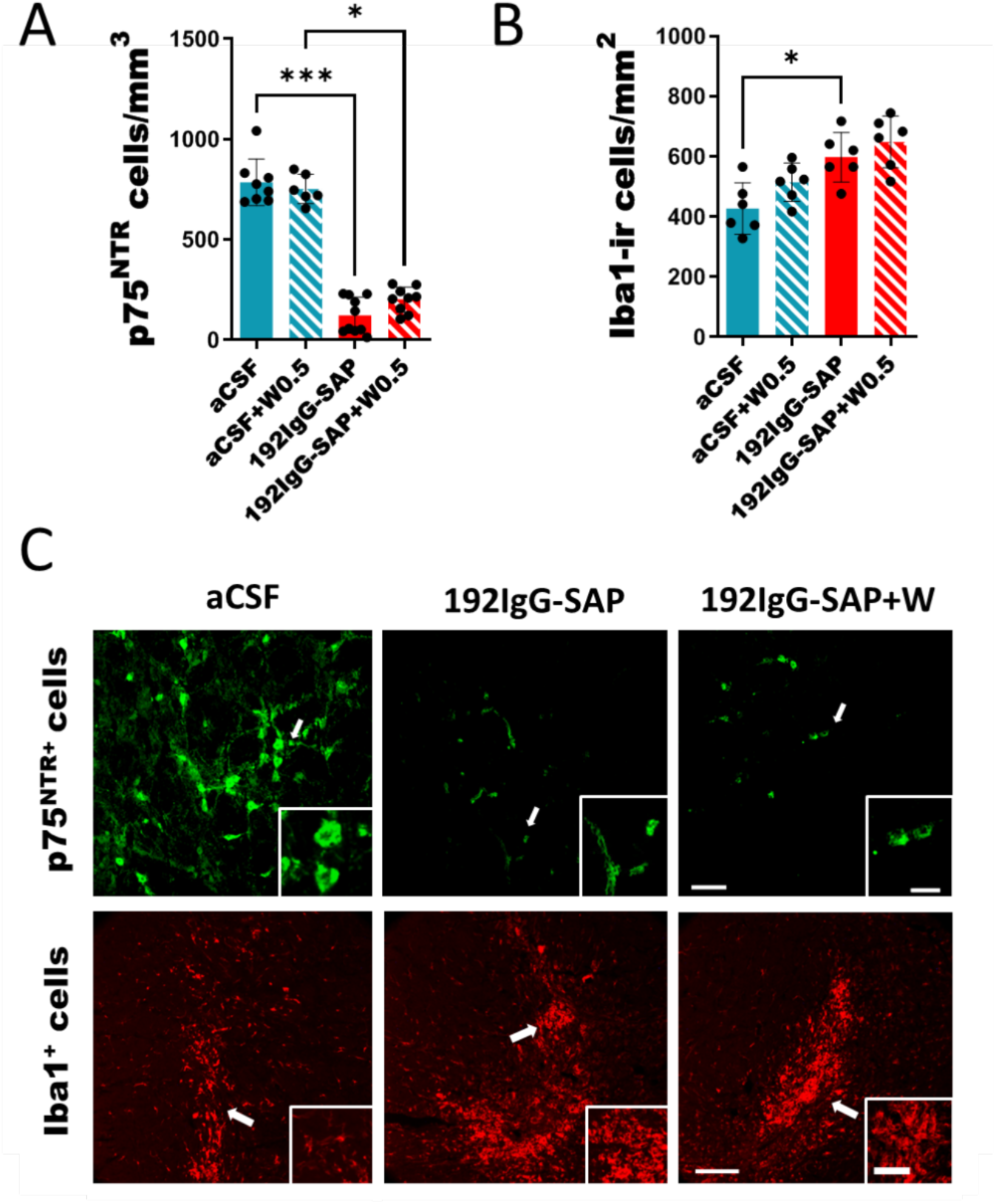
Immunofluorescent studies of (A) p75^NTR^ and (B) Iba1 positive cells of aCSF, 192IgG-SAP, aCSF +W and 192IgG-SAP+W groups in the NBM. (C) Labeling images of p75^NTR^ positive cells (green) and Iba1 positive cells (red) of aCSF, 192IgG-SAP and 192IgG-SAP+W (0.5 mg/kg) group. Note that 192IgG-SAP group had less p75^NTR^ positive cells (cholinergic cells) but had more Iba1 positive cells (microglia). WIN55,212-2 treatment did not modify cell number. Scale bar 100 µm (inset 50 µm) for p75^NTR^ and scale bar 100 µm (inset 25 µm) and Iba1 images. (Kruskal–Wallis test, *post-hoc* test Dunn’s multiple comparison \**p*<0.05 vs aCSF. Iba1: n=3; p75^NTR^: n=8).

Given the anti-inflammatory properties of cannabinoids and to describe potential changes in microglial phenotype or activation states^34^, microglial inflammation-associated lipid biomarkers in the same experimental groups were evaluated by MALDI-MSI (Supplementary Table 1). Following the lesion, a significant increase of lysophosphatidylcholines (LPC) C18 and C16 (LPC 18:0 a.u.; Control: 55207 ± 13922 *vs* 192IgG-SAP: 144700 ± 19651, LPC 16:0 a.u.; Control: 13808 ± 3085 *vs* 192IgG-SAP: 51672 ± 14782; \**p*<0.05), sphingomyelin (SM 34:1 a.u.; Control: 14531 ± 4591 *vs* 192IgG-SAP: 38556 ± 4956; \**p*<0.05) and palmitoyl (CAR 16:0) and oleoyl carnitine (CAR 18:1) levels were described at the lesion site (CAR 16:0; Control: 9893 ± 4149 *vs* 192IgG-SAP: 65156 ± 8637; CAR18:1; Control: 8120 ± 5517 *vs* 192IgG-SAP: 421720 ± 93521; \**p*<0.05), while C32-phosphatidylcholines (PC 32:0), showed a non-significant reduction. This lipidomic analysis showed that lipids associated to microglial inflammatory response increased at the lesion site. As an example, SM 34:1, which is restricted to the choroid plexus in physiological conditions, was detected at the lesion site following immunotoxin administration. Acyl-carnitines, which were only slightly detected in control rats, significantly increased in the lesion site following administration of the immunotoxin (Supplementary Figure 2). These results indicate that WIN55,212-2 administration did not restore these inflammation-associated lipids to control levels. Together with the results from the immunohistochemical studies showing unaltered number of microglial cells following the cannabinoid treatment, these results indicate that the subchronic administration of WIN55,212-2 did not significantly modify the post lesion microglia-associated inflammatory response in the NBM.

### WIN55,212-2 increased cortical muscarinic and cannabinoid receptor activity

The activity elicited by CB_1_ and muscarinic M_2_/M_4_ receptors (mAChR M_2_/M_4_) was analyzed in NBM, hippocampus and the cortex. The lesion selectively reduced the mAChR M_2_/M_4_ activity in layers III-IV of the cortex (mAChR M_2_/M_4_ of LIII-IV (% over the basal): aCSF: 350 ± 25; 192IgG-SAP: 105 ± 60; 192IgG-SAP+W: 242 ± 30, \**p*<0.05. Figure 3 A, C); with no observed effects in the NBM or hippocampus (Supplementary Figure 3). Conversely, low doses of WIN55,212-2 increased CB_1_ receptor activity in lesion animals in the same cortical layers ((% over the basal): 192IgG-SAP: 230 ± 70 vs 192IgG-SAP+W: 378 ± 21, ^#^*p*<0.05. Figure 3 A, B).

**Figure 3.**
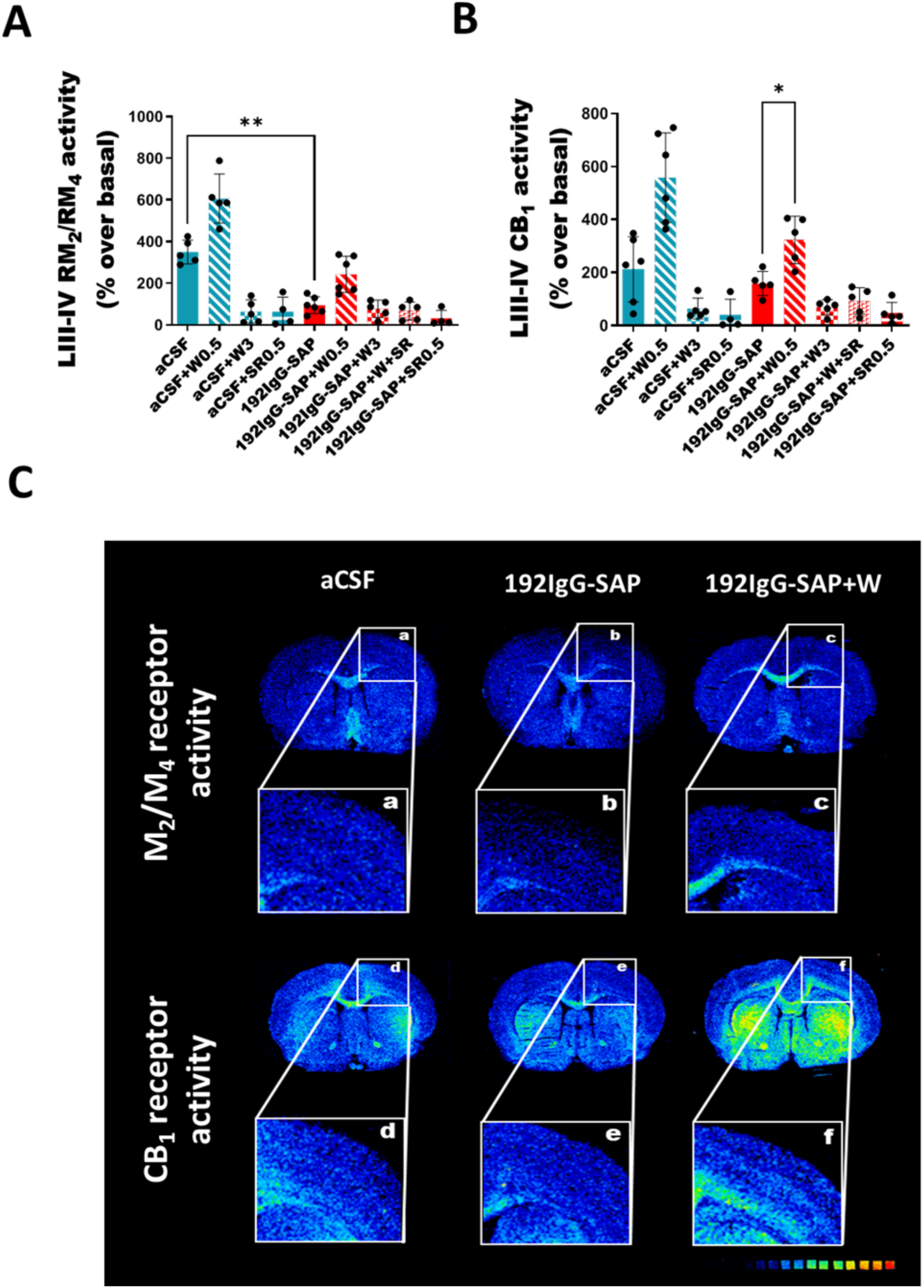
Functional autoradiographic studies of mAChR M_2_/M_4_ and CB_1_ receptor in cortical areas of all the experimental groups. **(A).** Graph of mAChR M_2_/M_4_ and CB_1_ receptor of all the experimental groups in layers III-IV of the cortex. Scale bar = 4 mm. (Kruskal–Wallis test, *post-hoc* test Dunn’s multiple comparison \*\*\**p*<0.001 **(B).** Representative autoradiographic images of brain coronal sections of mAChR M_2_/M_4_ and CB_1_R of aCSF, 192IgG-SAP and 192IgG-SAP+W0.5. Note that 192IgG-SAP+W0.5 group has same activity levels of mAChR M_2_/M_4_ and CB_1_ receptor than aCSF group.

### WIN55,212-2 modified cortical acetylcholinesterase activity, acetylcholine and choline levels

We measured acetylcholinesterase (AChE) activity, as well as ACh and choline (Ch) levels in the cortex in all the experimental groups. AChE activity in the cortex decreased after the lesion (AChE activity a.u.; aCSF: 17 ± 1 *vs* 192IgG-SAP: 8.4 ± 1, \**p*<0.001). Unexpectedly, the low/high doses of WIN55,212-2, as well as SR141716A, reversed the downregulation of AChE activity induced by the toxin (Figure 4 A, B).

**Figure 4.**
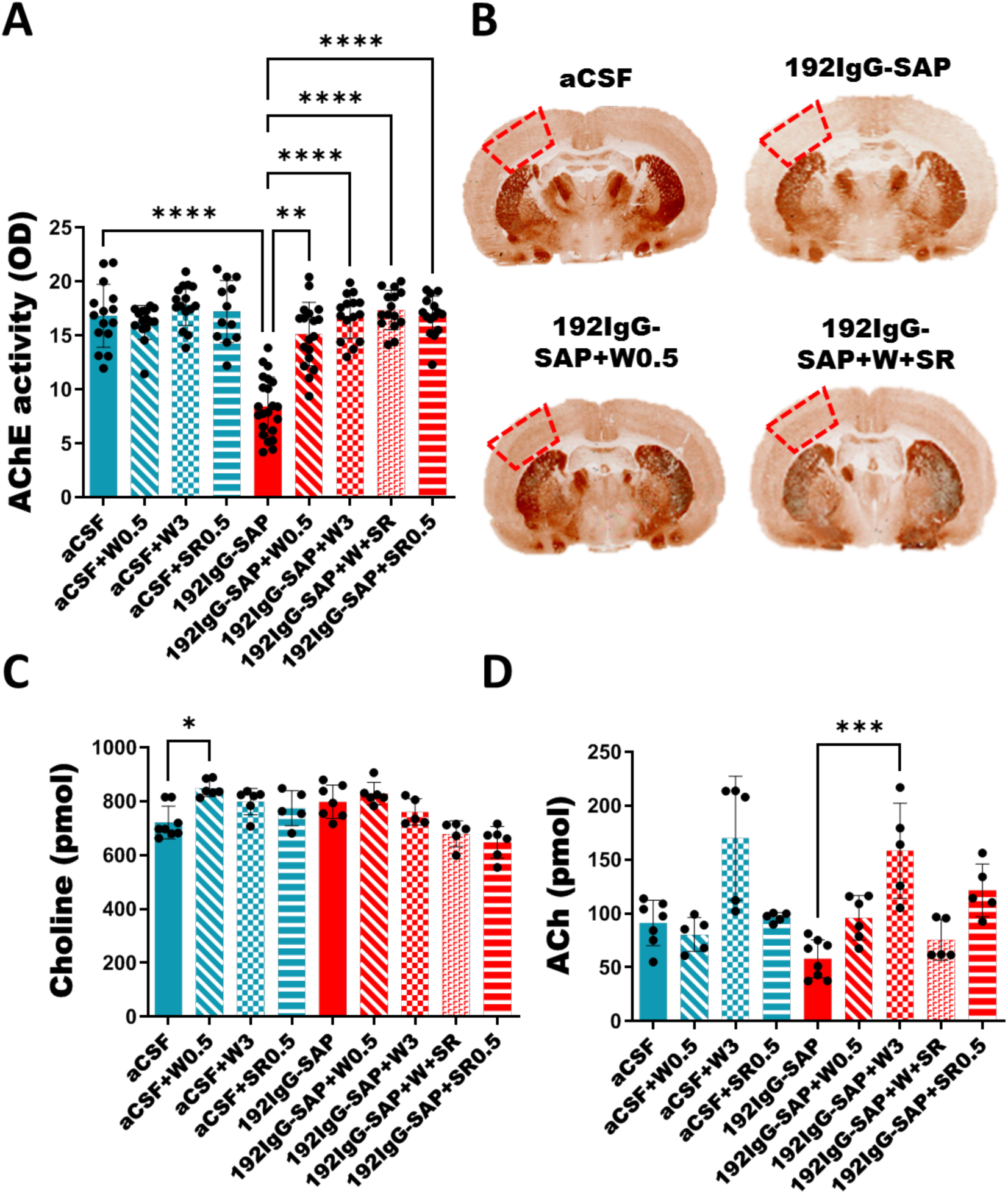
Cortical acetylcholinesterase (AChE) activity, choline (Ch) acetylcholine (ACh) and levels in all the experimental groups. (A). Boxplot of cortical AChE activity levels in all the experimental groups. (B). Representative images from coronal sections of AChE enzymatic staining from aCSF, 192IgG-SAP, 192IgG-SAP+W0.5 and 192IgG-SAP+W+SR groups. Note that WIN55,212-2 restores AChE cortical levels in the lesion animals (C). Boxplot of cortical choline levels in all the experimental group. (D). Boxplot of cortical choline levels in all the experimental groups. (Kruskal–Wallis test, *post-hoc* test Dunn’s multiple comparison. \*\*\**p*<0.001.)

Free cortical Ch levels (pmol) were increased in aCSF+W0.5 groups (aCSF: 721 ± 21 vs aCSF+W0.5: 798 ± 23, Figure 4 C) compared with the aCSF group. Additionally, cortical ACh levels showed a tendency to increase in the 192IgG-SAP+W0.5, but a significant increase was observed in the 192IgG-SAP+W3 group compared to 192IgG-SAP (192IgG-SAP: 58 ± 6, 192IgG-SAP+W0.5: 96 ± 8 and 192IgG-SAP+W0.5: 158 ± 19) (Figure 4 D). This pattern suggests a dose-response effect.

### WIN55,212-2 modified cortical lipid homeostasis

Cortical lipidomic analysis following WIN55,212-2 administration revealed more pronounced changes in lipid homeostasis compared to the impact of the cholinergic lesion alone (Figure 5 A). After the lesion, cortical saturated and mono-unsaturated lysophosphatidylcholine (LPC) levels (e.g., LPC 16:0 +K^+^, LPC 18:0 +K^+^ and LPC 18:1 +K^+^) were significantly reduced. There was also a reduction of two phosphatidylcholines (PC) (PC 38:2 and PC O-38:7) and an increase in phosphatidylethanolamine (PE) (PE 38:5).

**Figure 5.**
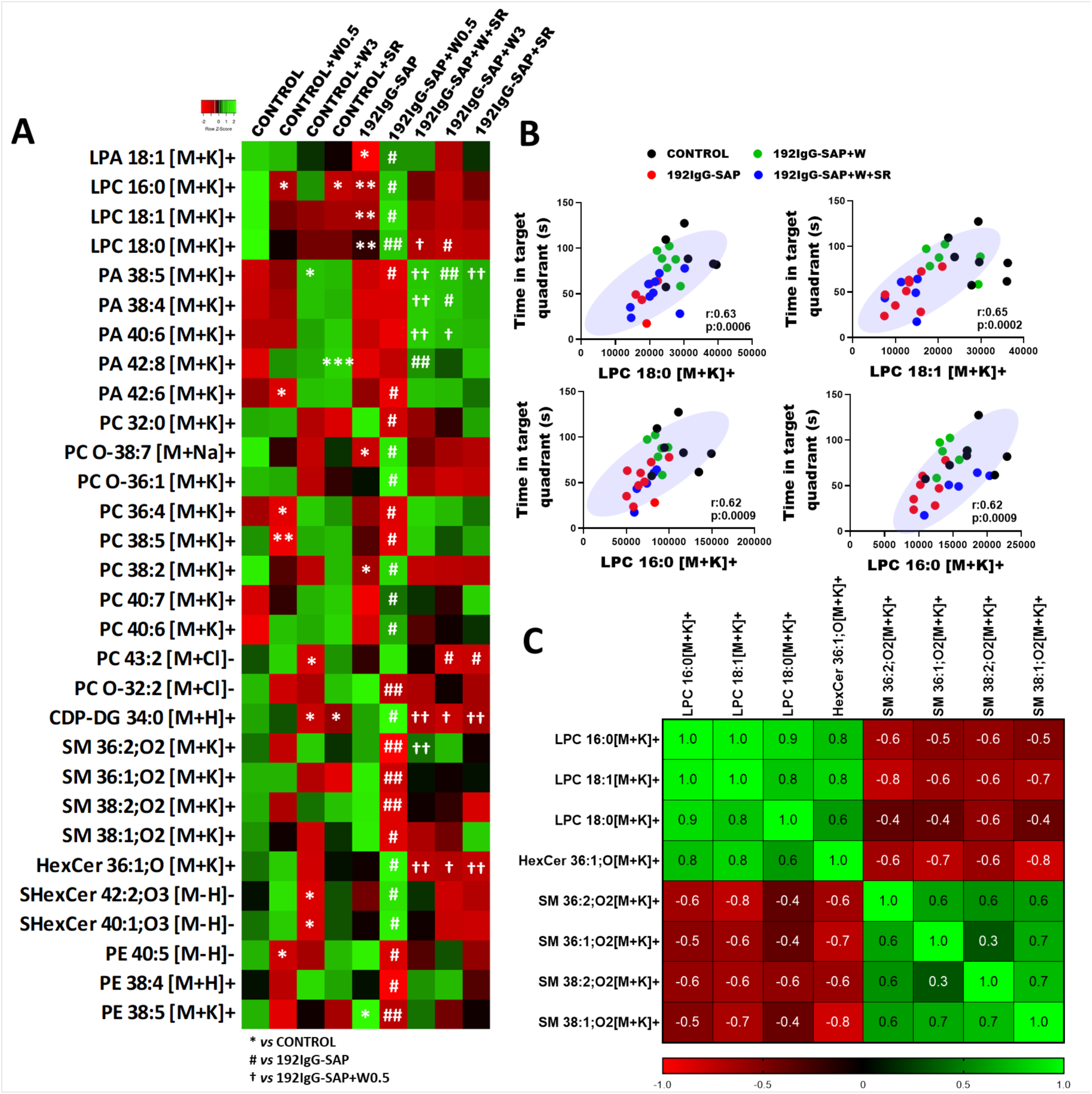
Cortical targeted lipidomic analysis. (A). Heatmap highlighting the 30 most differentially expressed lipid species between the different groups following the lesion or WIN55,212-2 treatment (Kruskal–Wallis test, *post-hoc* test Dunn’s multiple comparison \*\*\**p*<0.001). (B). Linear regression showing significant correlations between LPC 18:0, LPC 18:1, LPC 16:0 and LPA 18:0, and time in target quadrant on day 5 of Barnes maze (Spearman’s rank correlation coefficient r_s_ and p). (C). Matrix correlation between 192IgG-SAP and 192IgG-SAP+W, showing r_s_ values of Spearman’s correlations. Note the strong positive correlation between LPCs and HexCer, and the opposite correlation between LPC and SMs.

Lipidomic changes in 192IgG-SAP+W group, which displayed cognitive improvement after the lesion, revealed decreased arachidonic acid (AA) containing-phosphatidylcholines (PC) and phosphatidylethanolamines (PE) (e.g., PC 18:1_20:4 +K^+^, PC 16:0_20:4 +K^+^, PE 18:0_20:4 +H^+^, PE 18:1_20:4 +K^+^) and increased docosahexaenoic acid containing-phosphatidylcholines (DHA-PC), (e.g., PC 40:7 (18:1_22:6) +K^+^ and PC 40:6 (18:0_22:6) +K^+^). In addition, low doses of WIN55212-2 increased very long-chain sulfatides (e.g., SHexCer (d18:1_22:0)^-^, SHexCer (d18:1_24:1)^-^) and hexoceramides (HexCer 36:1 +K^+^) and decreased sphingomyelins (e.g., SM 36:1 +K^+^, SM 36:2 +K^+^, SM 38:1 +K^+^, SM 38:2 +K^+^). Furthermore, low doses of WIN55,212-2, increased LPCs to control levels. Interestingly, LPCs were the only lipids that correlated with behavioral parameters following BM testing in aCSF, 192IgG-SAP, 192IgG-SAP+W and 192IgG-SAP+W+SR groups, suggesting that LPCs are key players in the cognitive improvement seen in 192IgG-SAP+W animals (Figure 5 B). To identify the lipid precursor associated with LPC upregulation, correlations between lipid levels in 192IgG-SAP and 192IgG-SAP+W groups were performed. We found negative correlations with LPCs and SMs across groups, suggesting that increased LPCs are generated by SMs downstream pathway(s) (Figure 5 C).

### *In vitro* cortical SMs breakdown degradation produces increases in choline and LPCs

To better characterize the metabolic pathway linking LPCs, SMs, and choline, rat brain cortex tissue was incubated in a specific buffer to measure choline levels. Membrane lipids were then studied in the same sample after incubation using MALDI mass spectrometry. After a 2 h incubation, progressive diminution of some SM and increased levels of LPCs were found (Figure 6 A), as well as an increase in the choline levels (Figure 6 B). Notably, cortical tissue incubation replicated the same lipid changes observed after the WIN55,212-2 treatment *in vivo*. Here we report a strong correlation between ceramides/hexoceramides and choline levels, practically with a 1:1 ratio (Figure 6 C, D). SM degradation results in ceramide and phosphocholine, which is converted to choline (Figure 6 E), suggesting that the observed increase in choline found in the *in vitro* assay is related to the breakdown of SM. Moreover, LPCs increased conversely to SMs, in line with the observed changes seen after WIN55,212-2 treatment *in vivo*, further supporting the existence of this metabolic pathway which, in the *in vivo* model, is potentiated following CB_1_ receptor activation. To the best of our knowledge, our findings support for the first time that SM 36:1 is one of the alternative sources to produce choline, the precursor needed to produce the neurotransmitter acetylcholine and LPCs, respectively.

**Figure 6.**
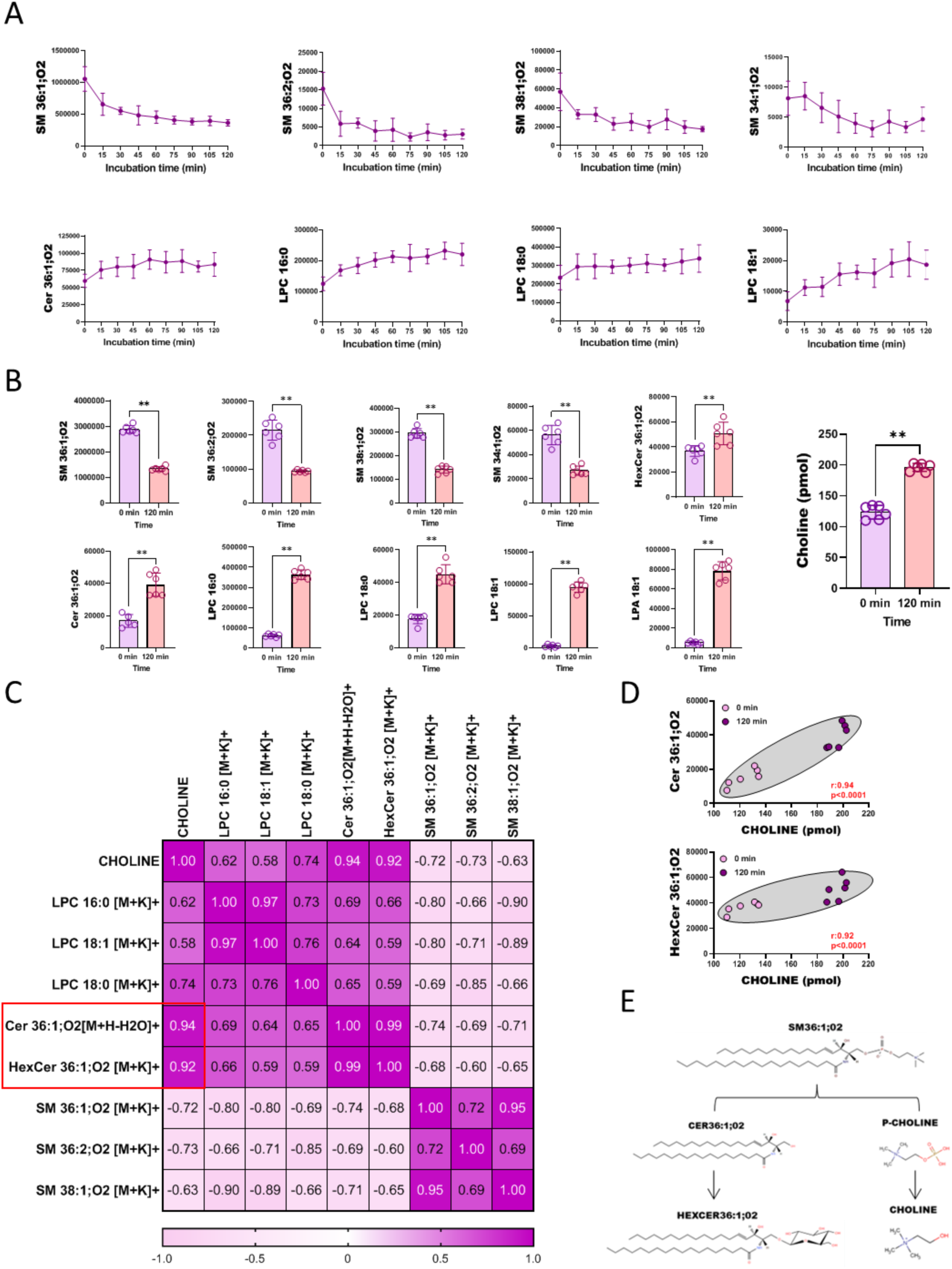
Choline production in brain cortex homogenates from control rats. **(A).** Line graphs of lipid production (LPCs) or degradation (SMs) in rat brain cortex homogenates over time (*n=8*, each time). **(B)**. Box plot of changes on cortical lipids and choline after 2 h of incubation (Mann-Whitney test, Time 0 *vs* Time 120 min, *n=6*). **(C)**. Correlation matrix between the LPCs, SMs and choline, showing r_s_ values of Spearman’s correlations, *n=6*. Note that choline and Cer/HexCer showed strong positive correlation. **(D)**. Linear regression shows significant correlations between Cer/HexCer and choline levels at time 0 and 120 min (Spearman’s rank correlation coefficient r_s_ and *p*). **(E)**. Degradation of sphingomyelins. Degradation of SMs generates ceramide and p-choline. Glycosylation of ceramide generates HexCer and dephosphorylation P-choline generates choline. Note that the same SMs and LPCs whose levels are modulated by WIN55,212-2 in the cortex of lesion rats change after incubation over time.

## DISCUSSION

The role that the eCB system plays in restoring cognitive impairment is an active area of research. Here we report *ex vivo* and *in vivo* data demonstrating that a low dose of the CB_1_ receptor agonist WIN55,212-2, improves cognition, following a lesion of the cholinergic neurons located in the NBM.

In the *ex vivo* study, application of 192IgG-saporin to P7 hemibrain organotypic cultures produced a significant cell death and specifically induced a loss of cholinergic neurons. Our results suggest that the application of the toxin triggers a partial loss of BFCN which leads to secondary cell damage, as demonstrated by the elevated levels of PI uptake. The established understanding is that p75^NTR^ facilitates the retrograde transport of both neurotrophins and the monoclonal antibody coupled to saporin from axon terminals^35^. This mechanism elucidates the depletion of BFCN subsequent to the uptake of saporin when administered intraventricularly *in vivo*^36^. In line with previous studies, the injection of the neurotoxin 192IgG saporin into the NBM *in vivo* resulted in a loss of p75^NTR^ neurons and cognitive impairment fifteen days post lesion^27, 28^. The administration of 192IgG-saporin produced a profound impairment of spatial, recognition and contextual memory in BM and NORT test, more pronounced in the long-term. The NBM primarily projects to the frontal, parietal, and temporal cortex. However, the tasks used here in BM and NORT also involve the hippocampus and associated cortical structures^37, 38^. We previously reported that no differences were observed in the hippocampus in terms of acetylcholinesterase (AChE) staining one week after the BFCN lesion^28^, providing evidence of the absence of nonspecific damage in other basal forebrain cholinergic projection pathways. This implies that the memory impairment observed in both BM and NORT mainly reflects the baso-cortical cholinergic damage.

Considering that the loss of cholinergic projections from the basal forebrain is an early pathological feature of clinical symptoms associated to dementia in AD^39, 40^, we extended our investigation to an animal model of familial AD, the 3xTg-AD mouse^30^. At 7 months, 3xTg-AD mice are at the onset of AD pathology, with cognitive functions already affected at this age^41, 42^. However, despite the moderate cognitive impact observed in 3xTg-AD mice at this stage, the BM, characterized as one of the most suitable tests for detecting cognitive deficits in this model at an early age^43^, did not reveal strong memory impairment. 3xTg-AD mice showed a prolonged learning curve as compared with matched wild-types, showing a delay in the spatial learning curve of this genotype at this age and no positive effects were observed following the treatment with the CB_1_ receptor agonist. Moreover, the time spent in the target quadrant, which is the main indicator to measure the acquired spatial memory in the test, was unaffected by genotype. Although the basal forebrain cholinergic system is affected early in this model^44^, it did not show significant memory deficits at least at this age and in this test. The focus of this study is to identify a treatment that specifically reverses memory impairment once severe cholinergic damage and memory deficits are already present. This makes the BFCN degeneration rat model a more valuable tool for investigating treatments against memory impairments induced by cholinergic deficits, thus, we decided to continue the neuropharmacological study using this rat model.

After successfully establishing our cholinergic lesion model *ex vivo* and also *in vivo* showing memory impairment after basal forebrain–cortical cholinergic circuits damage, we decided to initially test various doses of WIN55,212-2 in the *ex vivo* setting. The treatment of organotypic cultures with WIN55,212-2, pre-192IgG-saporin administration, showed protective effect against secondary cell death. Other authors previously demonstrated the WIN55,212-2 protective effect^45–47^. However, we found a non-significant impact specifically on the survival of cholinergic neurons. Nevertheless, those protective effects against secondary cell death *ex vivo* led us to administer the cannabinoid treatment in the *in vivo* lesion model following the occurrence of cholinergic cell loss. Thus, WIN55,212-2 would prove beneficial after the cholinergic damage has already taken place, i.e., when clinical cognitive symptoms arouse during neurodegenerative diseases.

The ip administration of a low dose (0.5 mg/kg) of WIN55,212-2 restored cognitive impairment on the *in vivo* model of BFCN degeneration on both BM and NORT. The recovery effect was mediated by CB_1_ receptors since co-administration of WIN55,212-2 with the specific CB_1_ receptor antagonist SR141716A blocked cognitive recovery. In addition, a high dose (3 mg/kg) of WIN55,212-2 had opposite effects depending on the treatment group including impaired memory in controls, while improving learning in lesioned animals in the BM. Similar effects were observed in NORT with the dose of 0.5 mg/kg. The administration of 0.5 mg/kg to the control rats impaired memory in the NORT, whereas it improved cognition in lesioned animals. It is widely accepted that cannabinoid agonism induces memory impairment, especially short-term memory^17, 48, 49^. Although numerous studies explore the role of cannabinoids in the impairment of spatial memory^50–54^, there are no studies conducted with cannabinoid agonism in rats using the BM. Meanwhile, studies in NORT test revealed that doses of WIN55,212-2 between 0.3-1.2 mg/kg are enough to completely impair short-term memory storage and different stages of long-term recognition memory^55, 56^. However, studies have reported either a biphasic effect of cannabinoid agonism on cognition^20^, or a beneficial effects of low as opposed to high doses^57, 58^. These results suggest that the dual effects of cannabinoids on cognition depend on several factors, more specifically, the status of the baso-cortical cholinergic pathway, as previously demonstrated by the cognitive status of the subjects^19^. Another important factor may be the inflammation status induced by intraparenchymal administration of 192IgG-saporin^59^, and the anti-inflammatory properties^60^of cannabinoids would be able to reduce it.

Although we found post toxin injection an increase in the number of Iba1 positive cells (microglia) in the region containing the NBM ^61^, the anti-inflammatory properties of WIN55,212-2^62, 63^ did not modify the number of microglia cells. Since there are multiple microglial phenotypes, it is difficult to perform an extensive analysis based on protein markers^64^. Instead, we used MALDI-MSI to identify inflammatory lipid patterns. Blank et al. ^34^, measured lipid profiles of lipopolysaccharide (LPS)-stimulated and unstimulated microglia-like cells and identified 21 potential inflammation-associated lipid markers. Inflammation-associated lipid markers studied by MALDI-MSI in the 192IgG group showed an increase in various lipids within the lesion site (e.g., LPC 18:0, LPC 16:0, SM 34:1 and palmitoyl/oleoyl-carnitines, CAR 16:0, CAR 18:1), but not in the control group. Following the ip administration of low doses of WIN55,212-2 in the 192IgG-SAP group, lipids did not decrease to control levels, indicating that WIN55,212-2 did not change either the number or the phenotype of microglia found at the lesion site. While several studies show that cannabinoid administration reduces inflammatory activity of microglia *in vitro* ^65–67^, no significant anti-inflammatory effect was observed following WIN55,212-2 administration with the dose and treatment protocol used in the present animal model. Considering the potential significance of the cholinergic system’s previous condition in the baso-cortical pathway for the positive impact of WIN55,212-2, and given the established interaction between the cholinergic and eCB systems^68, 69^, our study focused on analyzing how WIN55,212-2 affects or modulates the baso-cortical cholinergic pathway.

We found a decrease in cortical M_2_/M_4_ receptor activity and AChE activity after the lesion of the NBM. A similar impairment of muscarinic receptors has been described in AD, where there is a loss of cortical cholinergic innervation accompanied by a depletion of the M_2_ receptor^70, 71^. Low doses of WIN55,212-2 restored both cortical AChE and M_2_/M_4_ receptor activity. The lesion induced a non-significant trend towards reducing ACh levels, while low doses of WIN55,212-2 restored levels comparable to the control. In contrast, high doses of WIN55,212-2 significantly increased cortical ACh levels, surpassing those of the control group. In addition, WIN55,212-2 induced an increase in cortical free choline in control animals, where the cholinergic pathway was intact. It seems that, as mentioned earlier with the cognitive status, WIN55,212-2 also exhibit distinct effects on ACh and choline levels based on the state of the BFCNs. Thus, the cortical and subcortical ACh levels should be optimal for sustained attention, learning and memory^72^.

As for the eCB system, cortical CB_1_ receptor activity increased after WIN55,212-2 treatment in the lesioned animals. A comparable upregulation of the eCB system has been reported in early stages of patients with dementia^22^, indicating a compensatory response due to a loss of cholinergic innervation. This mechanism may play a role in the upregulation of cholinergic activity reported in the cortex of elderly people with mild cognitive impairment^73, 74^. We previously demonstrated an increase in cortical CB_1_ receptor activity following the degeneration of BFCN in response to cholinergic damage one week after the lesion^27^. However, our current findings indicate that two weeks after the lesion, there is no effect on the cortical CB_1_ receptor activity. During the progression of the lesion, the cannabinoid system primarily attempts to sustain balance by upregulating CB_1_ receptors, but it fails to maintain them over time. The baso-cortical cholinergic pathway may require continuous stimulation of cortical CB_1_ after the lesion to maintain proper memory functioning, as observed following the treatment with WIN55,212-2 in CB_1_ receptor cortical activity.

In this study, we showed that post-injury, despite the loss of the majority of BFCNs, WIN55,212-2 enhanced memory, likely by increasing cortical acetylcholine levels and maintaining the upregulation of CB_1_ receptor activity. Previous studies showed that cannabinoids modulate acetylcholine release in the hippocampus and cortex^24, 26, 75, 76^. However, the specific mechanisms underlying these effects remain incompletely understood. In this context, our study aimed to investigate the factors contributing to the increase in cortical ACh levels. Since cholinergic neurons use choline to synthesize ACh, but also to synthesize certain phospholipids, such as phosphatidylcholines and sphingomyelins^77^ we studied lipidomic changes following cannabinoid treatment using MALDI mass spectrometry.

Cortical lipidomic analysis revealed a significant decrease in some saturated and monounsaturated LPCs (LPC 18:0, LPC 18:1, and LPC 16:0) following the lesion. Our group had previously reported an increase in the production of these same LPCs with the activation of the muscarinic receptors^28^. Therefore, the reduction in cortical M_2_/M_4_ receptor activity after the lesion may impact the levels of these LPCs. These choline-containing compounds are being investigated for their potential as cognitive enhancers^11^ and play an inflammatory role due to their conversion to lysophosphatidic acid (LPA) and choline under the action of autotaxin^78^. These factors could offer alternative explanations for the reduction observed after the BFCN lesion; hence, maintaining the homeostasis of these molecules may be crucial for memory. Surprisingly, low doses of WIN55,212-2 restored these cortical LPCs to control levels, correlating with cognitive scores obtained in the BM. This supports the importance of maintaining cortical homeostasis for these molecules, although the mechanism of recovery for these LPCs was unknow.

To identify the lipidic changes induced by WIN55,212-2 associated with the restoration of cortical LPCs, correlations were performed between lipidomic data derived from the 192IgG-SAP group (showing decreased LPC levels) and the 192IgG-SAP+W group (where LPC levels were restored). Although low doses of WIN55,212-2 induced numerous changes in cortical lipidomic homeostasis, the most significant alterations were the decreases in SMs (SM 36:1, SM 36:2, SM 38:1, SM 38:2), which correlated with the increased LPCs. The negative correlation observed between this type of lipids *in vivo* may suggest that SMs degradation induces the increment of LPCs. Choline plays a crucial role as a key metabolite necessary to maintain homeostasis between these two lipids^79^. Both LPCs and SMs are choline-containing lipids, although they do not directly share metabolic pathways^80,81^. In this sense, the selective vulnerability of BFCNs in various neurodegenerative diseases might result, in part, from an imbalance between these choline-containing lipids, as choline is used to synthesize both choline-containing lipids and ACh.

To better understand the homeostasis of these lipids and their relationship with choline, we incubated naive cortical rat brain tissue in specific buffers for two hours to measure choline levels and lipid composition *in vitro* within the same sample. The assay revealed that the increase in choline was accompanied by elevated LPCs and Cer/HexCer, as well as a decrease in SMs, similar changes to those observed after WIN55,212-2 administration *in vivo*. The strong correlation observed between choline and Cer/HexCer *in vitro* indicates that choline is specifically derived from the breakdown of cortical SMs. Sphingomyelinases, both acid and neutral, directly act by cleaving the phosphocholine group, transforming SM into ceramide^82^. Phosphocholine is a derivative of choline, which is crucial for the synthesis of ACh^83^. These results indicate that low doses of WIN55,212-2, through SM degradation, generate an alternative cortical choline source, as observed in control rats, used to increase both ACh and LPC levels. Similarly, the generation of ACh through PC decomposition has been described, but not through SM degradation^76^. This newly reported source of choline may originate from astrocytes, given that these SMs (SM 36:1, SM 36:2, SM 38:1, SM 38:2) are predominantly present in astrocytic membranes^84,85^, supporting earlier reports demonstrating cannabinoid-mediated breakdown of SMs in astrocytic cultures^86–88^. Collectively, these findings support our hypothesis regarding the memory enhancement mechanism in lesioned rats. Following BFCN degeneration, WIN55,212-2, through the continuous activation of CB_1_ receptors in the cortex (possibly in astrocytes), triggers sphingomyelinase activation. This process generates ceramide and phosphocholine, ultimately producing free choline. Consequently, cortical levels of acetylcholine and LPCs experience an increase. This is reflected in restored AChE activity, M_2_/M_4_ receptor activity, and their correlation with cognitive improvement.

The present findings indicate the need to maintain cortical choline-containing lipids, for the normal functioning of the cholinergic basal forebrain cortical projection system and cognition. Therefore, pharmacological modulation of the eCB system may represent a promising therapy for neurodegenerative disorders involving basal forebrain cholinergic degeneration, such as AD.

## ACKNOWLEDGEMENTS

Technical and human support provided by University of the Basque Country (UPV/EHU), Ministry of Economy and Competitiveness (MINECO), Basque Government (GV/EJ), European Regional Development Fund (ERDF), and European Social Fund (ESF) is gratefully acknowledged. J.M.-G. is the recipient of “Margarita Salas” fellowship and I. B de T. of an “Investigo” fellowship funded by the European Union-Next Generation EU. The authors would also like to show gratitude to Dr. Elliott J. Mufson and Dr. Sylvia E. Perez from the Barrow Neurological Institute for their assistance with paper revisions and editing.

## AUTHOR CONTRIBUTIONS

M.M.-R and R.R.-P. conceived and designed the study, performed the statistical analysis, and wrote the manuscript. A.L.-O and L.L. contributed to organotypic cultures studies, such as developing an *ex vivo* model of cholinergic lesion and *in vitro* treatments. J.M.-G., M.M.-R. and E.G.d.S.R. contributed to the performance of MALDI-MSI experimental procedures and data analysis. M.M.-R, J.M.-G and I.B.d.T. performed the *in vivo* studies, such as surgeries, treatments, and behavioral tests. M.M.-R, J.M.-G and I.M. performed autoradiographic studies. M.M.-R performed acetylcholinesterase activity, acetylcholine, and choline assays. M.M.-R and J.M.-G. performed the lipidomic analysis in incubated and non-incubated samples by MALDI. All authors contributed to the manuscript revision, read, and approved the submitted version.

## Data and material availability

The data that support the findings of this study are available from the corresponding author upon reasonable request. Some data may not be made available because of privacy or ethical restrictions.

---

**Supplementary Figure 1.**
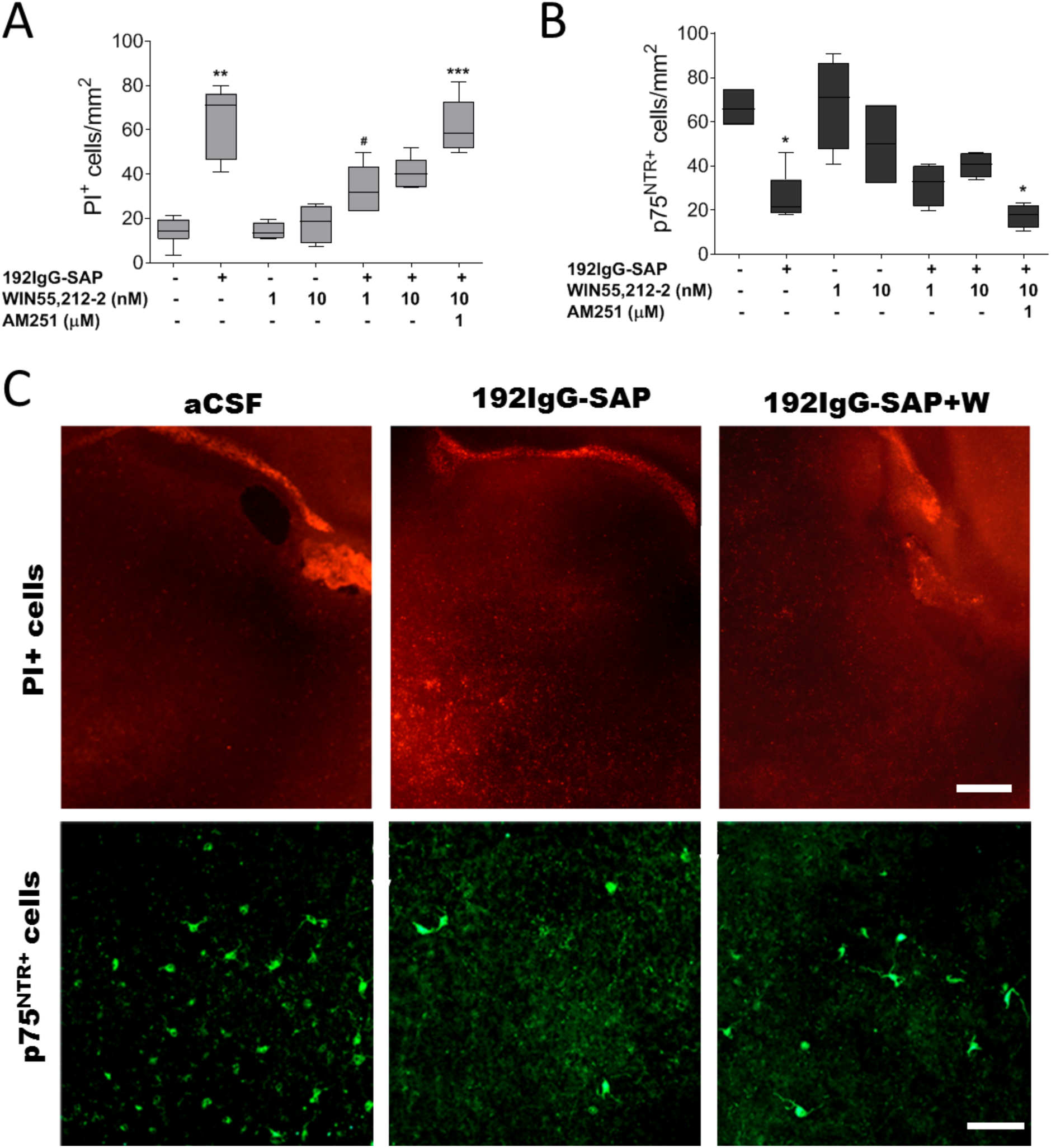
*Ex vivo* model of organotypic culture treated with 192IgG-saporin and WIN55,212-2. **(A).** Number of PI^+^ cells and **(B).** p75^NTR+^ cells in organotypic cultures in the absence or presence of 100 ng/ml of 192 IgG-saporin, cannabinoid agonist WIN55,212-2 (1 or 10 nM) and antagonist AM251 (1 µM) in *NBM* (Kruskal-Wallis test, *post-hoc* test Dunn’s multiple comparison **\*\****p*<0.01 *vs* CONTROL, ^#^*p*<0.05 *vs* 192IgG-SAP. **PI:** *n*=7 CONTROL; *n*=4 C+W(1nM); *n*=4 C+W(10nM); *n*=6 192IgG-SAP; *n*=6 192IgG-SAP+W(1nM); *n*=5 192IgG-SAP+W(10nM); *n*=6 192IgG-SAP+W+AM251. **p75^NTR^:** *n*=5 CONTROL; *n*=4 C+W(1nM); *n*=4 C+W(10nM); *n*=4 192IgG-SAP; *n*=4 192IgG-SAP+W(1nM); *n*=4 192IgG-SAP+W(10nM); *n*=4 192IgG-SAP+W+AM251. **(C).** Representative images of PI^+^ cells (red) and p75^NTR+^ (green) immunoreactivity in NBM of control, 192IgG-SAP and 192IgG-SAP+W(1nM). PI^+^ cell Scale bar = 200 µm and p75^NTR^ scale bar = 40 µm.

**Supplementary Figure 2.**
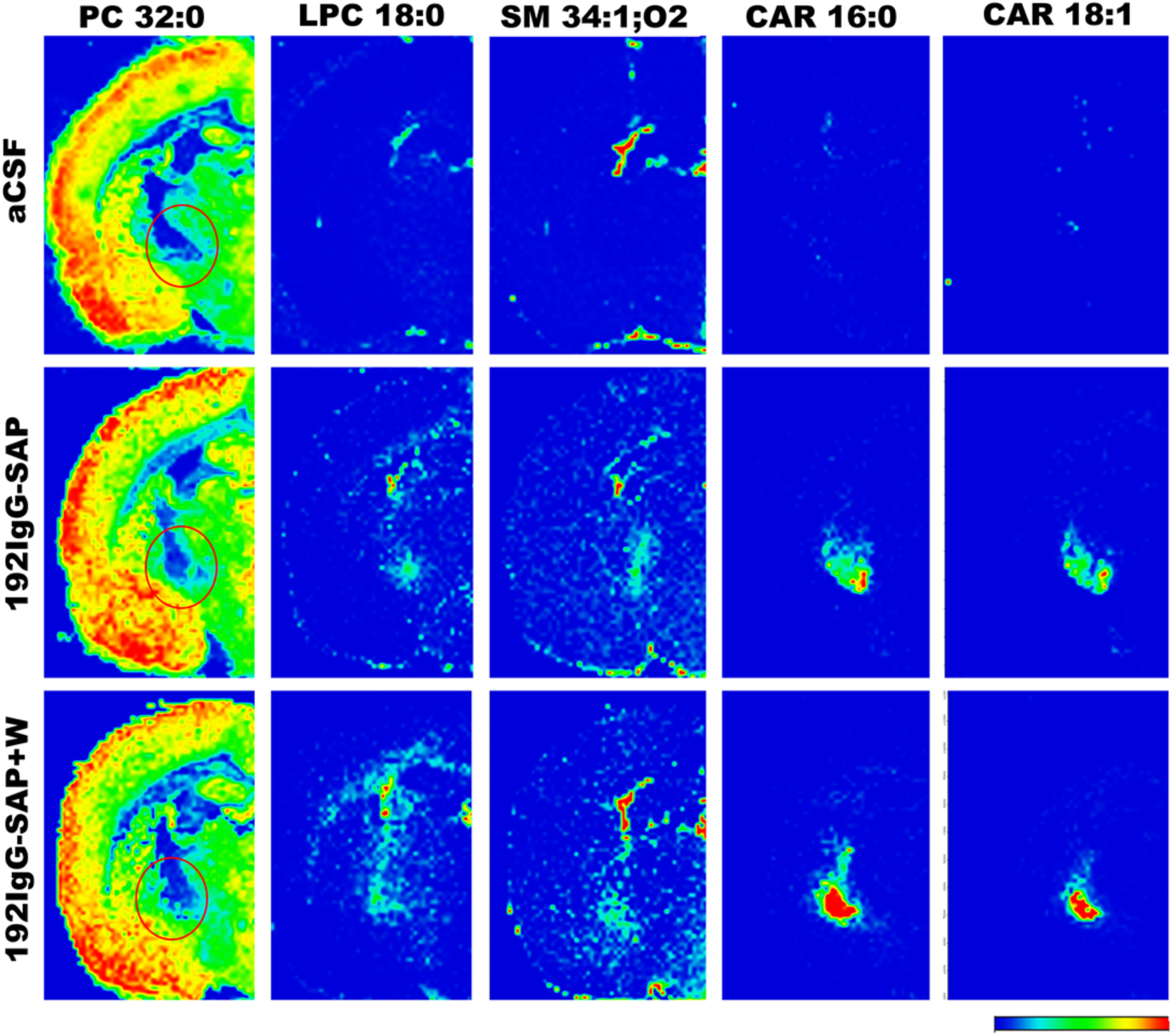
Matrix-assisted laser desorption ionization mass spectrometry imaging (MALDI-MSI) of inflammation-associated lipids in brain coronal slices of Control, 192IgG-SAP and 192IgG-SAP+W0.5 group. Red circles indicate the lesion site, where lipids related to inflammation-associated microglia were measured. Note that lysophosphatidylcholine (LPC 18:0), sphingomyelin (SM 34:1), palmitoylcarnitine (CAR 16:0) and oleoylcarnitine (CAR 18:1) levels increased specifically at the lesion site. *n*=5.

**Supplementary Figure 3.**
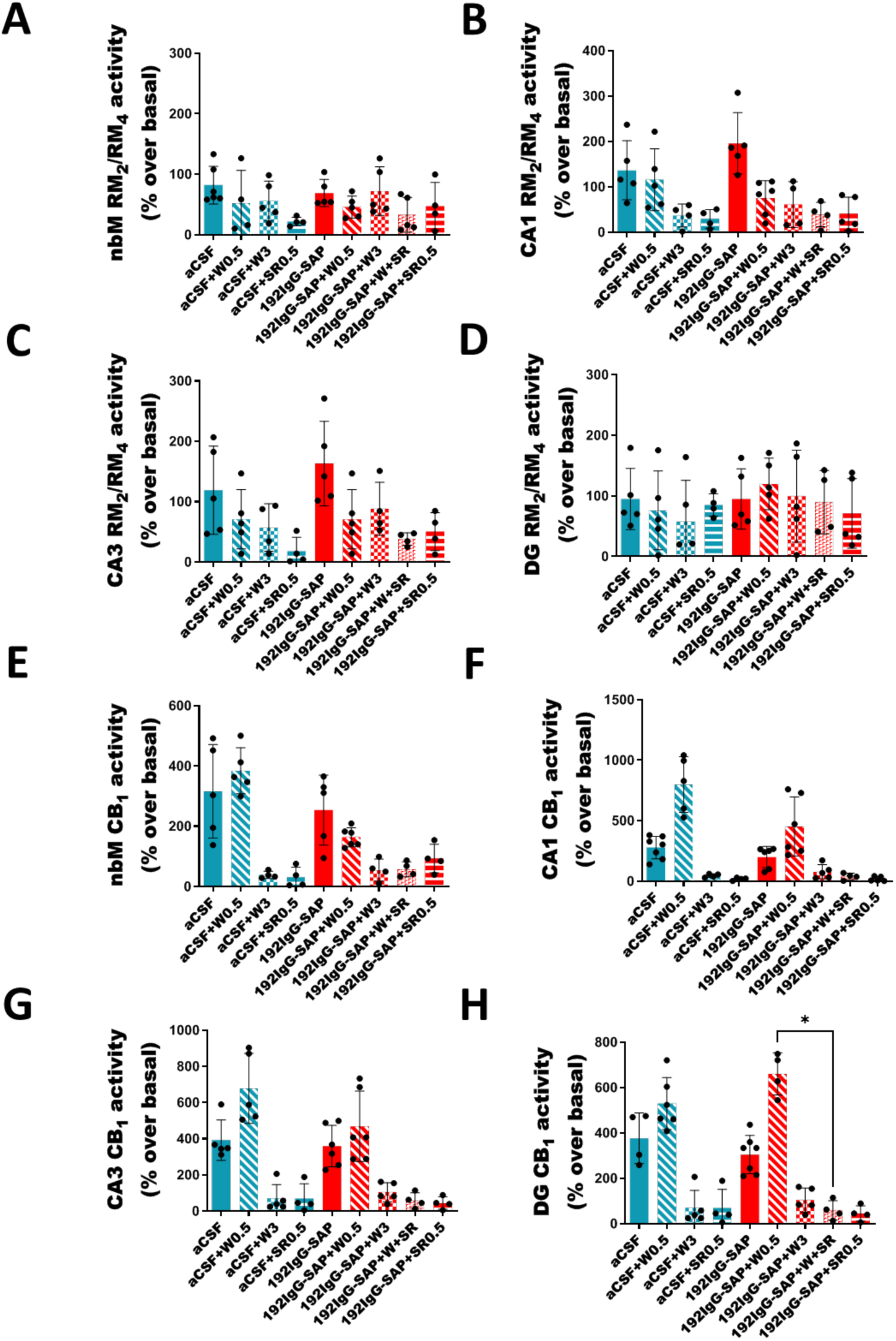
Functional autoradiographic studies of mAChR M_2_/M_4_ and CB_1_ receptors in NBM and hippocampal areas of all the experimental groups. (A) mAChR M_2_/M_4_ activity in the NBM (B) mAChR M_2_/M_4_ activity in the CA3 of the hippocampus. (C) mAChR M_2_/M_4_ activity in the CA1 of the hippocampus. (D) mAChR M_2_/M_4_ activity in the dentate gyrus (DG) of the hippocampus. (E) CB_1_ receptor activity in the NBM (F) CB_1_ receptor activity in the CA3 of the hippocampus. (G) CB_1_ receptor activity in the CA1 of the hippocampus. (H) CB_1_ receptor activity in the dentate gyrus (DG) of the hippocampus. Note that there are no significant differences between the aCSF and 192IgG-SAP groups, as well as between the 192IgG-SAP and 192IgG-SAP+W0.5 groups.

**Table 1.** Absolute intensity of inflammation-associated lipid species biomarkers in microglia in the lesion site after low doses of WIN55,212-2 in positive mode.

| Assignment | Cal m/z | Exp m/z | CONTROL | 192IGg-SAP | CONTROL+W0.5 | 192IGg-SAP+W0.5 |
| --- | --- | --- | --- | --- | --- | --- |
| LPC O-16:0[M+H]<sup>+</sup> | 482.3605 | 482.3623 | 2545 ± 970 | 5232 ± 2204 | 6583 ± 2720 | 9267 ± 3043 |
| LPC 16:0 [M+H]<sup>+</sup> | 496.3398 | 496.3415 | 213258 ± 12803 | 296546 ± 27758 | 189738 ± 23642 | 245438 ± 30677 |
| LPC 16:0 [M+Na]<sup>+</sup> | 518.3217 | 518.3237 | 2373 ± 875 | 3989 ± 1528 | 3291 ± 762 | 4313 ± 1014 |
| LPC 18:0 [M+H]<sup>+</sup> | 524.3711 | 524.3730 | 55207 ± 13922 | 144700 ± 19651* | 83098 ± 13201 | 132257 ± 10238 |
| LPC 16:0 [M+K]<sup>+</sup> | 534.2956 | 534.2978 | 13808 ± 3085 | 51672 ± 14782* | 34680 ± 11128 | 57686 ± 13633 |
| LPC 18:0 [M+K]<sup>+</sup> | 562.3278 | 562.3289 | 3486 ± 1312 | 24647 ± 4448* | 16286 ± 7514 | 23697 ± 4108 |
| DG 34:0 [M+H-H<sub>2</sub>O]<sup>+</sup> | 579.5347 | 579.5345 | 75434 ± 28780 | 54819 ± 15955 | 47509 ± 15491 | 55641 ± 18476 |
| Cer 42:1;O<sub>2</sub> [M+H-H<sub>2</sub>O]<sup>+</sup> | 632.6340 | 632.6358 | 5793 ± 2008 | 5870 ± 1717 | 14170 ± 1850* | 8458 ± 1933 |
| PC 32:0 [M+H]<sup>+</sup> | 734.5694 | 734.5722 | 2.07*10<sup>6</sup> ± 475451 | 730389 ± 206873 | 1.07*10<sup>6</sup> ± 269945 | 979923 ± 208596 |
| PC 32:0 [M+Na]<sup>+</sup> | 756.5513 | 756.5543 | 912569 ± 279206 | 842267 ± 79401 | 586950 ± 111616 | 592423 ± 64619 |
| PC 32:0 [M+K]<sup>+</sup> | 772.5253 | 772.5278 | 3.32*10<sup>6</sup> ± 499928 | 2.59*10<sup>6</sup> ± 213702 | 2.66*10<sup>6</sup> ± 270507 | 2.54*10<sup>6</sup> ± 247160 |
| PC 34:0 [M+K]<sup>+</sup> | 800.5578 | 800.5566 | 488070 ± 102509 | 525862 ± 148953 | 521543 ± 107415 | 556673 ± 107250 |
| SM 34:1;O<sub>2</sub> [M+K]<sup>+</sup> | 741.5307 | 741.5333 | 14531 ± 4591 | 38556 ± 4956* | 20205 ± 3815 | 34127 ± 8424 |
| CAR 16:0 [M+H]<sup>+</sup> | 400.3421 | 400.3434 | 9893 ± 4149 | 65156 ± 8637* | 102026 ± 30789 | 424809 ± 59210# |
| CAR 18:1 [M+H]<sup>+</sup> | 426.3578 | 426.3585 | 8120 ± 5517 | 57604 ± 15119* | 65605 ± 34259 | 421720 ± 93521# |

Data are mean ± S.E.M values of absolute intensity of Control (n = 5), 192IGg-SAP (n = 5), Control+W0.5 (n=5), 192IGg-SAP+W0.5 (n = 5) groups. Kruskal–Wallis test, *post-hoc* test Dunn's multiple comparison \**p*<0.05 vs Control, #*p*<0.05 vs 192IgG-SAP. PC: phosphatidylcholine; DG: diacylglycerol, SM: sphingomyelin, CER: ceramide, LPC: phosphatidylcholine and CAR: carnitine; Cal: calculated; Exp: experiment

